# Development of a universal nanobody-binding Fab module for fiducial-assisted cryo-EM studies of membrane proteins

**DOI:** 10.1101/2021.08.20.457137

**Authors:** Joël S. Bloch, Somnath Mukherjee, Julia Kowal, Ekaterina V. Filippova, Martina Niederer, Els Pardon, Jan Steyaert, Anthony A. Kossiakoff, Kaspar P. Locher

## Abstract

With conformation-specific nanobodies being used for a wide range of structural, biochemical, and cell biological applications, there is a demand for antigen-binding fragments (Fabs) that specifically and tightly bind these nanobodies without disturbing the nanobody-target protein interaction. Here we describe the development of a synthetic Fab (termed NabFab) that binds the scaffold of an alpaca-derived nanobody with picomolar affinity. We demonstrate that upon CDR grafting onto this parent nanobody scaffold, nanobodies recognizing diverse target proteins and derived from llama or camel can cross-react with NabFab without loss of affinity. Using NabFab as a fiducial and size enhancer (50 kDa), we determined the high-resolution cryo-EM structures of nanobody-bound VcNorM and ScaDMT, both small membrane proteins of ~50 kDa. Using an additional anti-Fab nanobody further facillitated reliable initial 3D structure determination from small cryo-EM test datasets. Given that NabFab is of synthetic origin, humanized, and can be conveniently expressed in *E. coli* in large amounts, it may not only be useful for structural biology, but also for biomedical applications.

## Introduction

Antibodies have long been a cornerstone of cell biological and biochemical research due to their unique ability to bind with exquisite selectivity to an immense range of molecular targets. Engineering antibody fragments to introduce new functionalities is made possible by their modular form, which combines a conserved fold with a set of hypervariable loops (complementary-determining regions, CDRs) that confer the antibody’s specificity and binding affinity. This conserved organization has guided endeavors to engineer novel forms and formats with enhanced properties to fabricate antibody-like molecules with increased functionality as research tools and biotherapeutic entities(1–7).

Antibodies have also been beneficial for structural biology. However, raising monoclonal antibodies using traditional hybridoma methodology has often constituted a significant barrier. In recent years, engineered antibody fragments have had a major impact on structural biology, particularly by accelerating the structure determination of membrane proteins. They were either applied as crystallization chaperones(8) or selected to trap specific states of a target protein, thereby reducing conformational flexibility. While the recent emergence of electron cryomicroscopy (cryo-EM) has altered the structure determination landscape significantly, determining high-resolution structures of small particles (currently ~50 kDa) or highly dynamic macromolecules including membrane proteins remains challenging(9). Two classes of antibody fragments have been widely and successfully used for structure determination: (i) Fab (fragment antigen binding) domains that consist of two protein chains and have an approximate molecular mass of ~ 50 kDa and (ii) heavy chain-only antibody fragments, also known as nanobodies (Nbs), which are found in camelids and sharks(5) and have an approximate molecular mass of ~ 14 kDa. Unlike Fabs, Nbs are devoid of light chains and the CDRs are located on a single Ig domain of the heavy chain (VHH). Due to their groove- or cavity-binding propensity, Nbs can target epitopes that are inaccessible to the larger Fabs. This has facilitated the trapping and stabilization of membrane proteins such as transporters, ion channels and receptors in distinct conformational states(10–14).

We and others have shown that Fab fragments are a game changer for cryo-EM studies of smaller proteins, particularly membrane proteins where the surrounding detergent micelles or lipid nanodiscs impair the ability to align particles(15–20). While individually generating Fabs against membrane protein targets is possible, a more general approach using a universal fiducial would be desirable. In previous manifestations of universal fiducials, an “off-the-shelf” Fab was generated to a binding motif that was introduced into the protein under investigation. For instance, we have designed and implemented universal Fab based fiducials to a BRIL fusion(21, 22), to different classes of trimeric G-proteins(23, 24), and to portable stem-loop RNA motifs(25). Additionally, a variety of other types of portable motifs have been developed(26, 27), some of which involve nanobodies applied as the portable motif in a different context(28).

Given that many nanobodies have been raised to trap or induce specific conformational states of target proteins, a universally applicable anti-Nb Fab would be beneficial. In this embodiment, the Nb provides the function, the Fab provides the size and shape. We therefore pursued a strategy to develop a “universal” fiducial that combines the attributes of both the Nb and Fab fragments into one unit. We describe here the development and implementation of a universal anti-Nb Fab (NabFab) that can be readily applied to virtually any nanobody-protein complex. NabFab has been engineered to attach distal to the Nb’s CDR loops and at an angle that ensures that it does not interfere with the nanobody’s target. We show how the NabFab is compatible with virtually all Nb scaffolds, enabling easy plug and play use. We demonstrate the utility of NabFab by determining high resolution cryo-EM structures of two distinct ~50 kDa membrane proteins. Capitalizing on the humanized scaffold(29, 30) of the NabFab, we employed a second nanobody that binds to the hinge connecting the variable and constant domains of the NabFab light chain(31). This adds a distinct feature that breaks the pseudo-symmetry of the Fab to facilitate the analysis of three-dimensional reconstructions in small cryoEM test datasets. With these findings, we demonstrate the applicability of NabFab as a powerful fiducial for cryo-EM thereby enabling the structure determination of many important biological systems that have existing nanobody binders available to them. To facilitate broad use of NabFab, the expression plasmid and associated protocols are freely available upon request.

## Results and Discussion

### Generation and characterization of a synthetic anti-nanobody Fab (NabFab)-

The workflow for developing, validating, and utilizing a universal nanobody-Fab system is outlined in Figure 1a. In a first step, a cohort of anti-Nb Fabs was generated by phage display mutagenesis utilizing a high-diversity (~10^10^) synthetic phage display library based on a humanized antibody Fab scaffold(29). The target antigen was an alpaca-derived Nb (TC-Nb4) against the complex of human transcobalamin and its cognate receptor TCblR/CD320(32). To obtain Fabs that bind an epitope distal to the CDRs of TC-Nb4, the selection was performed in the presence of molar excess of its cognate target TC:CD320(33), which effectively masks the CDR loops. After four rounds of selection, the enrichment of specific binders over non-specific background clones was >50 fold, and DNA sequencing of clones that gave a strong signal over background in single point phage ELISA, resulted in 12 unique binders. The high diversity in CDR-H3 and L3 indicated that the cohort of clones represented a large diverse pool. Eleven of the unique clones were sub-cloned in the Fab format and successfully expressed in *E. coli* and purified in large scale.

**Fig. 1.**
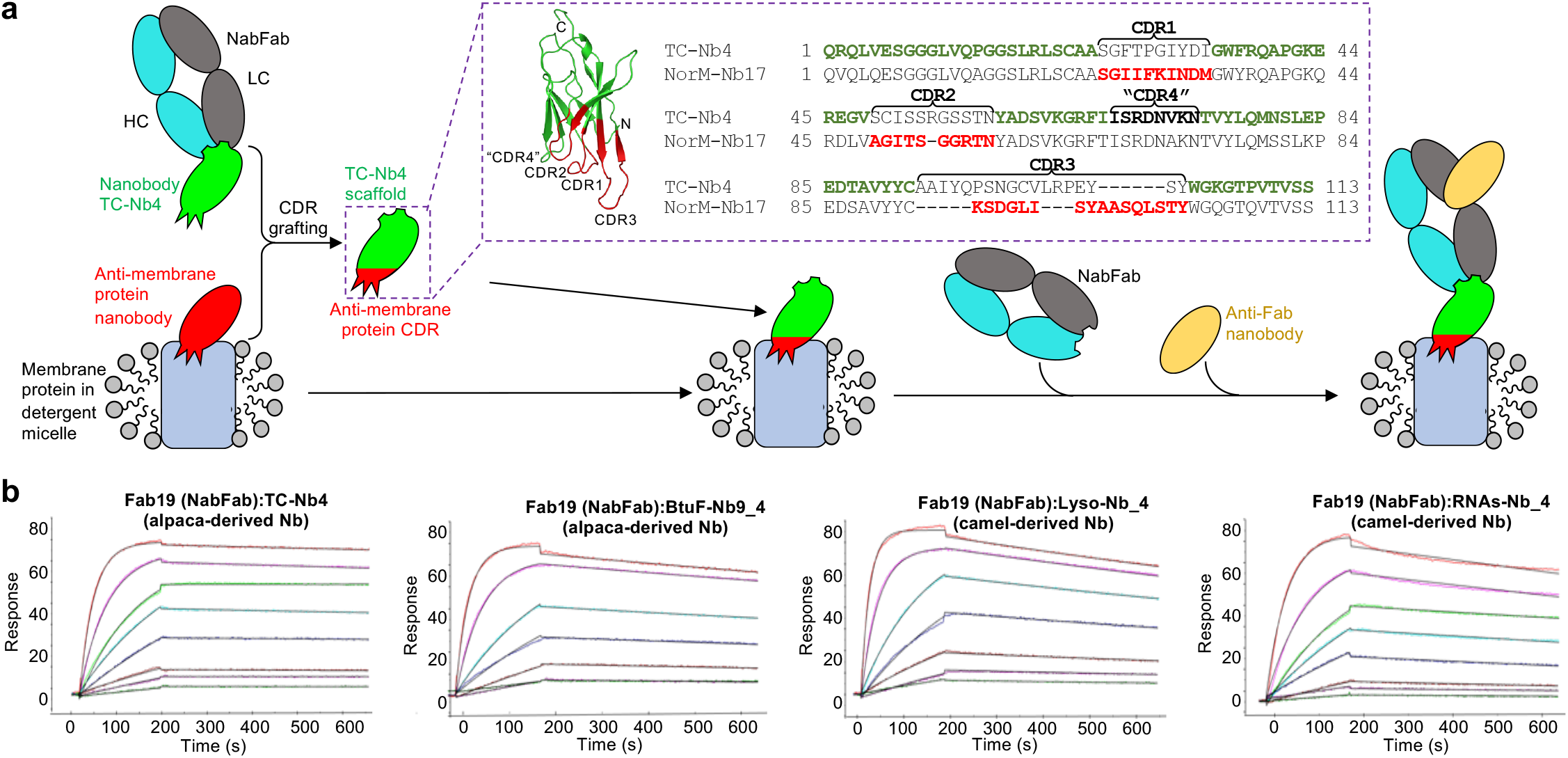
Universal binding of NabFab to the scaffold nanobodies enabled by CRD grafting. **a,** Schematic of NabFab application in Nb binding. Grafting of target-specific Nb CDRs onto the TC-Nb4(32) scaffold, which contains the NabFab binding epitope, enables rigid complex formation of target membrane protein, grafted Nb, NabFab, as well as an anti-Fab Nb(31) as symmetry breaker. Dashed boxes illustrate the rationale behind CDR grafting. On the example of the structure of NorM-Nb17_4 (ribbon representation) the TC-Nb4 scaffold(32) is shown in green and the grafted CDRs of NorM-Nb17 in red. On the right side a sequence alignment of TC-Nb4 and NorM-Nb17 is shown in Kabat numbering(47). The scaffold and CDR regions that comprise the grafted NorM-Nb17_4 are highlighted with the same colors. Note that for NorM-Nb17_4 “CDR4” of TC-Nb4 was retained due to its similarity in sequence. **b**, SPR sensograms of NabFab binding to TC-Nb4 (alpaca) and grafted nanobodies BtuF-Nb9_4 (alpaca), Lyso-Nb_4 (camel), and RNAs-Nb_4 (camel).

While Nbs have a common structural scaffold, their surface residues are not fully conserved. We anticipated that the Fabs selected against TC-Nb4 might bind an epitope that was not fully compatible with all possible Nb scaffold sequences. We therefore adopted a grafting strategy to ensure the binding of other Nbs, no matter their sequence variation or CDR loop composition, to the Fab selected against TC-Nb4. The CDR loops from target Nbs were inserted in the scaffold of TC-Nb4 to obtain chimeras that retain the binding properties of the original Nb. Conceptually, this is the same process involved in the “humanization” of antibodies. We tested the feasibility of the approach using Nbs of distinct origin by swapping their CDR loops into the scaffold of TC-Nb4 guided by the boundaries show in Fig. 1a. We used Nbs targeting the vitamin B12-binding bacterial protein BtuF(34), lysozyme(35), and RNAseA(36) resulting in the chimeras BtuF-Nb9_4, Lyso-Nb_4, and RNAs-Nb_4. Analytical size exclusion chromatography combined with gel electrophoresis experiments (Supplementary Data Fig. 2) demonstrated that the chimeric Nbs bound their cognate antigen targets, indicating the robust nature of the engineered scaffold. Next the cross-reactivity of the 11 generated Fabs to the three Nb chimeras was tested by a single-point protein ELISA experiment (Supplementary Data Fig. 1). While all Fabs bound to the parent TC-Nb4 scaffold, several showed little or no cross-reactivity to the Nb chimeras. We speculate that the binding of these Fabs may be affected by the antigen used in the selection, demonstrating the importance of experimentally confirming that the Fabs to be carried forward are devoid of antigen influence. Fab14 and Fab19 displayed strong binding to all four chimeric Nb constructs and the binding affinity and kinetics to TC-Nb4 were measured by surface plasmon resonance (SPR). Fab19 and Fab14 bind to TC-Nb4 with affinities (KD) of 860 pM and 21 nM, respectively (Table 1). While both Fab14 and Fab19 are high affinity binders for most structural biology applications, Fab19 is superior because it has a ~60-fold slower off-rate than Fab14. We evaluated the binding of Fab19 to the other three chimeric Nbs, demonstrating that it bound all three with low nM affinity (Table 1, Fig. 1b).

**Table1.** Kinetic parameters of Fabs binding to wt TC-Nb4 and scaffold grafted nanobodies.

| Fab | Nanobody | $k_{on}$ ($M^{-1}s^{-1}$) | $k_{off}$ ($s^{-1}$) | $K_D$ (nM) |
| --- | --- | --- | --- | --- |
| Fab14 | TC-Nb4 | $4.2 \times 10^5$ | $8.8 \times 10^{-3}$ | 21 |
| NabFab (Fab19) | TC-Nb4 | $1.6 \times 10^5$ | $1.4 \times 10^{-4}$ | 0.9 |
| | BtuF-Nb9_4 | $1.1 \times 10^5$ | $4.2 \times 10^{-4}$ | 3.9 |
| | Lyso-Nb_4 | $2.2 \times 10^5$ | $3.5 \times 10^{-4}$ | 1.6 |
| | RNAs-Nb_4 | $1.8 \times 10^6$ | $3.0 \times 10^{-4}$ | 1.7 |

We next evaluated how Fab19 would perform in the context of Nbs binding to membrane proteins. We selected three Nbs targeting the bacterial MATE transporter VcNorM (llama-derived NorM-Nb17_4), the bacterial divalent metal ion transporter ScaDMT(37) (llama-derived DMT-Nb16_4) and the bacterial oligosaccharyltransferase PglB(38) (alpaca-derived PglB-Nb17_4). As in the case with the soluble targets, SEC analysis demonstrated that Fab19 binds to the chimeric Nbs in the context of their cognate membrane protein targets (Supplementary Data Fig. 2). Thus, based on Fab19's high affinity, superior kinetic profile, and its performance with chimeric Nbs derived from different species and targeting both soluble and membrane protein systems, it was designated as the lead candidate (NabFab) to be further tested as a universal nanobody fiducial in a series of cryo-EM experiments.

### Fiducial-Assisted single particle cryo-EM structure determination

To assess the potential of NabFab as a fiducial to enhance size and shape to assist in alignment of small particles for cryo-EM structure determination, we chose two ~50 kDa-sized membrane proteins, VcNorM and ScaDMT. These membrane proteins have no significant extra-membranous features and are of a size that is representative of many important biotherapeutic targets and remain challenging targets for cryo-EM. While the X-ray structure of VcNorM has been reported without bound nanobodies(39), that of ScaDMT had been determined in complex with a llama-derived nanobody (DMT-Nb16)(37). For VcNorM we generated a specific nanobody by immunization of llama(40). For the use with NabFab, we generated the chimeric nanobodies NorM-Nb17_4 and DMT-Nb16_4.

VcNorM and ScaDMT were purified in detergent (dodecyl maltoside, DDM) and incubated with the grafted chimeric Nbs, NabFab, and an anti-Fab nanobody that had been raised to bind to the elbow linker between the variable and constant domains of the Fab's light chain (31). In our previous work, we found this anti-Fab Nb markedly improved the attributes of the Fab fiducials. First it reduces the inherent flexibility between the variable and constant domains, which is Fab-dependent(41). Second, even though the size of the elbow nanobody only marginally increases the mass of the NabFab, it adds a distinctive element to the shape of the fiducial that assists in particle alignment. Combining the four components yielded stochiometric complexes (Supplementary Data Fig. 2) that were directly applied to cryo-EM grids and subjected to cryo-EM analysis (Fig. 2). Note that NabFab does not bind the scaffold of the anti-Fab nanobody, which prevents Fab-nanobody polymerization.

**Fig. 2.**
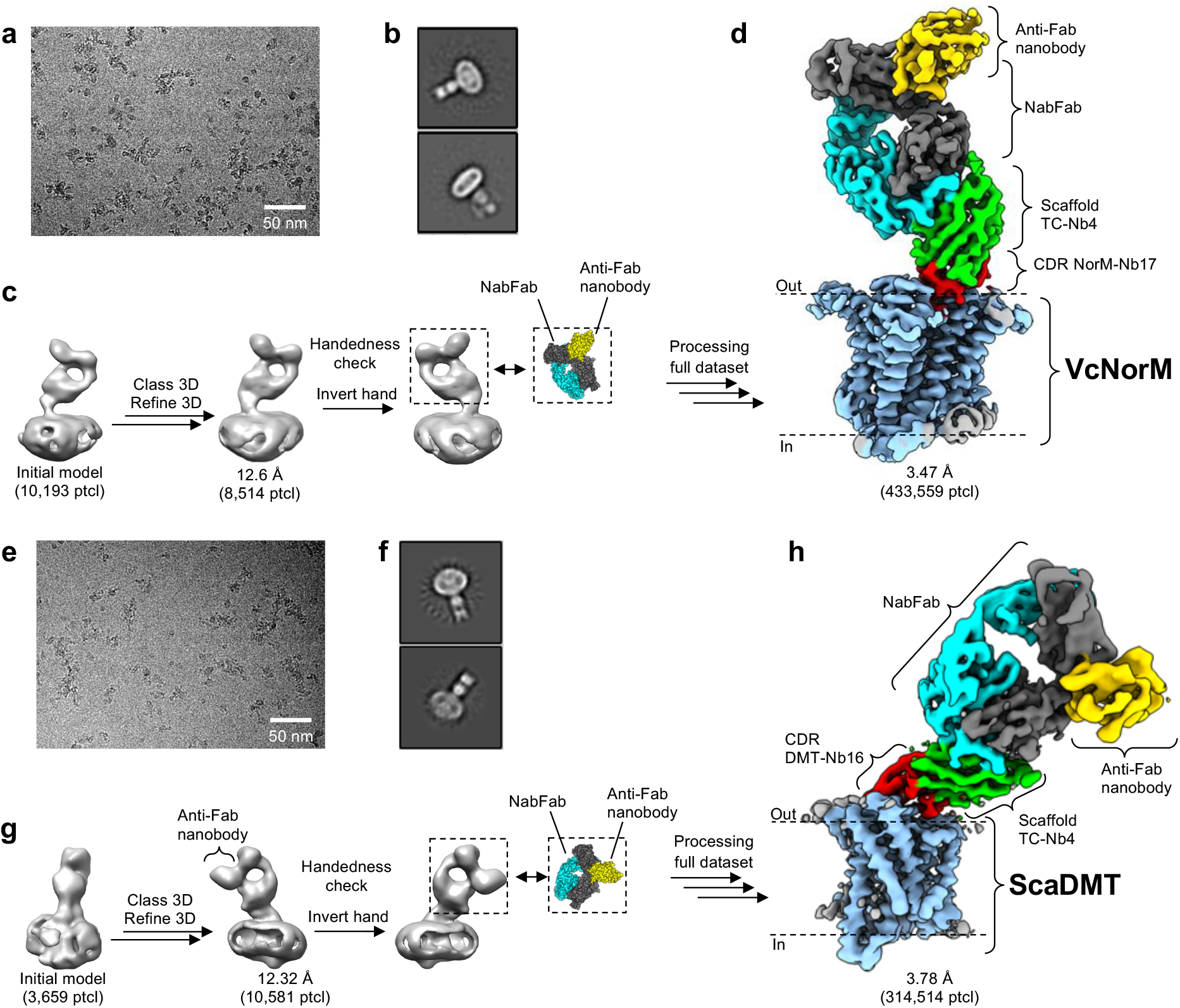
Fiducial assisted structure determination. **a-d,** Cryo-EM structure determination of VcNorM complex. **e-h,** Cryo-EM Structure determination of ScaDMT complex. **a/e**, Representative cryo-EM micrographs of complexes of detergent-solubilized membrane protein, grafted Nb, NabFab, and anti-Fab Nb. **b/f**, Representative 2D classes as calculated in RELION 3.1(42)**. c/g**, Processing of a small cryoEM test dataset including initial model generation and 3D refinement as calculated in RELION 3.1(42) and assignment of the correct handedness of the structure using structural knowledge of NabFab:anti-Fab Nb complex. **d/h**, Unsharpened Cryo-EM maps of VcNorM and ScaDMT complexes derived from the processing of full datasets (VcNorM in RELION 3.1 and ScaDMT in RELION 3.1(42) and cryoSPARC V3.2(48)). VcNorM and ScaDMT are highlighted in light blue, grafted Nb CDRs in red, TCNb4 scaffold in green, NabFab HC in cyan and LC in black, and the anti-Fab Nb in yellow.

The distinct structural features of the complexes were readily visible in the motion-corrected micrographs (Fig. 2 a, e) and became more evident during 2D classification (Fig. 2b, f). We were able to determine initial models from sets of particles obtained from 380 (VcNorM) or 625 (ScaDMT) micrographs that were collected in under 2 hours at a Titan Krios cryo-TEM. For each complex, a reconstruction could be refined to ~12 Å resolution, using as little as ~10,000 particles (Fig. 2c, g). At this stage of analysis, two features of the fiducial are readily apparent. First, there is a discernable hole between the variable and constant domains of NabFab, which is a hallmark of all Fab fiducials. Second, the anti-Fab Nb provides a distinct feature that allows the handedness of the obtained reconstruction to be correctly assigned even at low resolution given the characteristic shape of the NabFab:anti-Fab Nb complex(31). This is advantageous for rapid analysis of small cryo-EM test datasets using NabFab.

Upon collecting larger datasets, we obtained 3D reconstructions at final resolutions of 3.47 Å for VcNorM and 3.78 Å for ScaDMT (Fig. 2d, h, Supplementary Data Figs. 3, 4, Supplementary Data Table 3). In both structures, a rigid NabFab-Nb interface was observed (Supplementary Data Figs. 5, 6). Local resolution estimates indicate that differences in the flexibility of the two membrane proteins might have led to the observed resolution differences. The higher rigidity of VcNorM compared to ScaDMT can probably be attributed to the bound Nb NorM-Nb17_4, whose CDRs are deeply wedged into the outward-open cavity of the transporter. In the VcNorM structure, we increased the local resolution of different regions of the complex by shifting of the map fulcrum (Supplementary Data Fig. 5), which was originally placed at the center of the membrane protein by the processing software RELION3.1(42). VcNorM was in a similar outward-facing state as in previously published structures(39). However, the improved resolution allowed us to build a *de novo* model, which revealed that significant registry errors exist in the published structure. A detailed analysis of the cryo-EM structure of VcNorM will be published elsewhere.

For ScaDMT, our EM map was of similar quality as that of the previously published X-ray structures(37). A difference we observed was the dimerization state of ScaDMT with DMT-Nb16. While the original DMT-binding Nb promotes dimerization of the DMT-Nb complex in solution and in the crystal lattice(37), the chimeric DMT-Nb16_4 does not allow a similar association of DMT-Nb complexes. We therefore observed mainly monomeric complexes of DMT and no nanobody-mediated dimerization. However, we identified a subclass of ScaDMT-Nb complex dimers (Supplementary Data Fig. 4b, f) with a distinct arrangement from the published structure(37). Although the resolution of this dimeric reconstruction was low (~8.5 Å), the chiral NabFab-anti-Fab nanobody-handle allowed us to unambiguously fit ScaDMT into the EM density. The observed dimer interface is formed by TM7 and EL3-H1 of ScaDMT and does not appear to be caused or influenced by any of the involved Fabs or nanobodies. This finding may reflect a mode of homodimerization for ScaDMT or other SLC11/NRAMP family members.

### Fab nanobody interface

In addition to the two cryo-EM structures, we also determined a 3.2 Å crystal structure of NabFab in complex with Lyso-Nb_4 (Supplementary Data Table 4) and compared it with the cryo-EM structures containing VcNorM and ScaDMT. While the three complexes each contained a single copy of NabFab and a bound Nb, the CDRs of the Nbs are distinct. Remarkably, the three NabFab-Nb interfaces (Fig. 3a-d) are indistinguishable at the present resolution, demonstrating the effectiveness of our CDR grafting strategy. An interface analysis revealed the conserved interactions among the three structures (Supplementary Data Table 2). The two EM maps and the X-ray map are well-resolved at the NabFab-Nb interface and even revealed bound water molecules, most of which are visible in all three structures (Fig. 3e-g). This underscores the structural rigidity of the interface, which is an important factor to consider in determining the potential of a fiducial mark.

**Fig. 3.**
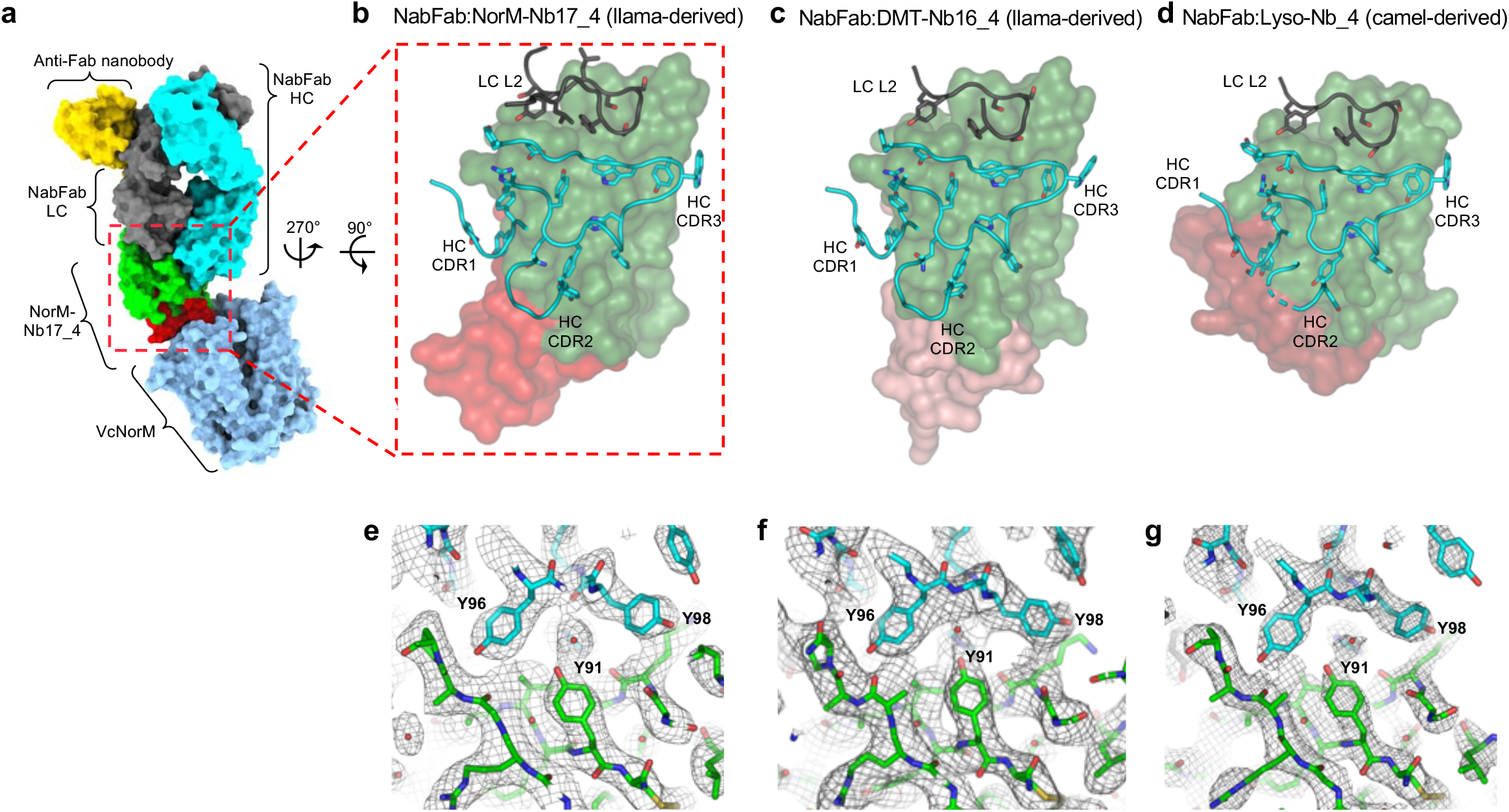
Structural analysis of NabFab binding to grafted nanobodies of different origin. **a**, Surface representation of complex of VcNorM (blue), NorM-Nb17_4 (TC-Nb4(32) scaffold green, NorM-Nb17 CDRs red), NabFab (HC cyan, LC black), and anti-Fab Nb(31) (yellow). **b**, Zoomed-in view of NorM-Nb17_4:NabFab binding interface with NorM-Nb17_4 in surface representation and NabFab in ribbon representation with interface residues in stick representation with oxygen atoms colored in red and nitrogen atoms in blue. **c**, DMT-Nb17_4:NabFab binding interface, **e**, Lyso-Nb_4:NabFab binding interface **d**, Representative section of EM map of NorM-Nb17_4:NabFab interface at 12 rmsd, carved to 2 Å. Ordered water molecules indicated by red crosses. **e**, Representative section of EM map of DMT-Nb16_4:NabFab interface at 11 rmsd, carved to 2 Å. **f**, Representative section of electron density of Lyso-Nb_4:NabFab interface at 1 rmsd, carved to 2 Å.

NabFab binds the β-strands of the TC-Nb4 scaffold mostly via its long CDR3 loop of the heavy chain (HC). Additional contacts are provided by the CDR1 of the HC and by the CDR2 of the light chain (LC). The C-terminal His-tags of the NorM-Nb17_4 and DMT-Nb16_4 nanobodies are partially ordered in the EM density maps and appear to weakly interact with the LC scaffold, but given the absence of a His-tag in purified Lyso-Nb_4 molecule, these contacts are likely not essential. The buried surface area between NabFab and the Nb scaffold amounts to an average of 976 A^2^ (731 ± 23 A^2^ for the HC and 245 ± 18 Å^2^ for the LC). There are several prominent π-π and cation-π interactions involving aromatic NabFab residues (Fig. 4a). To identify the hotspot residues that must be retained during scaffold grafting, we generated alanine mutants of the interface residues of the CDR-grafted LysoNb_4 nanobody and tested their respective affinities to NabFab by SPR (Fig. 4b, Supplementary Data Table 1). The most significant reduction in affinity was observed for P41 and V89, where mutations to alanine reduced the affinity between 400-600-fold, resulting in ΔΔG° values of 3.6 and 3.8 kcalmol^−1^, respectively (Figs. 4b, 5a). Substantial effects were also observed upon mutation of R45 (>40-fold higher KD) and Y91 (>25-fold higher KD), whereas mutating L11, K43, W103, or S112 to alanine only marginally (<5-fold) reduced the affinity of NabFab to LysoNb_4. Notably, our alignments suggest that NabFab binding extends beyond camelid-derived nanobodies and, based on sequence alignments, appears to be compatible with the scaffold used in the synthetic nanobody libraries NbLib(43) and Sybody(44) (Supplementary Data Fig. 7).

**Fig. 4.**
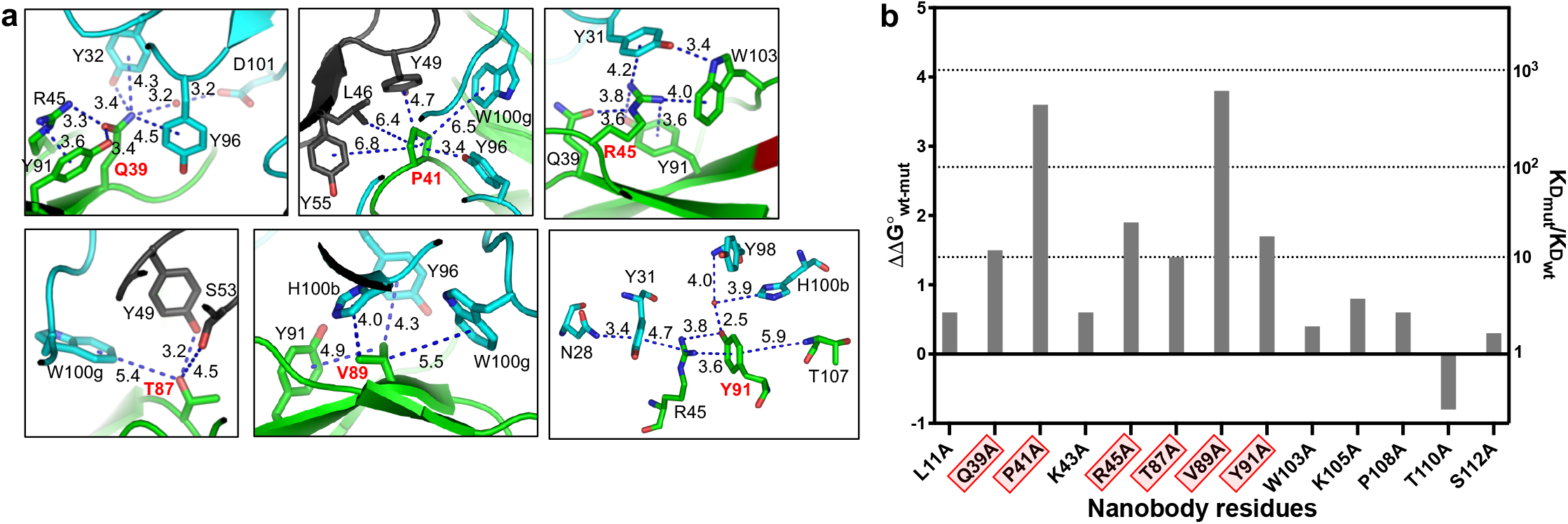
Hotspot analysis of NabFab binding epitope on TC-Nb4 scaffold. **a**, Detailed views of NorM-Nb17_4 residues at the NabFab binding interface. Distances in Å are indicated with numbers and dotted lines. Note that R45 and Y91 are contributing to a cation-π ladder. **b**, Alanine scanning of nanobody residues at the interface to NabFab. Bars indicate the ratio of the K_D_ for NabFab binding to the original TCNb4 scaffold and the respective surface mutants as derived from SPR measurements. Residues for which mutation to Ala reduced binding affinity more than 10-fold are highlighted in red. Note that Lyso-Nb_4 was used in this experiment.

**Fig. 5.**
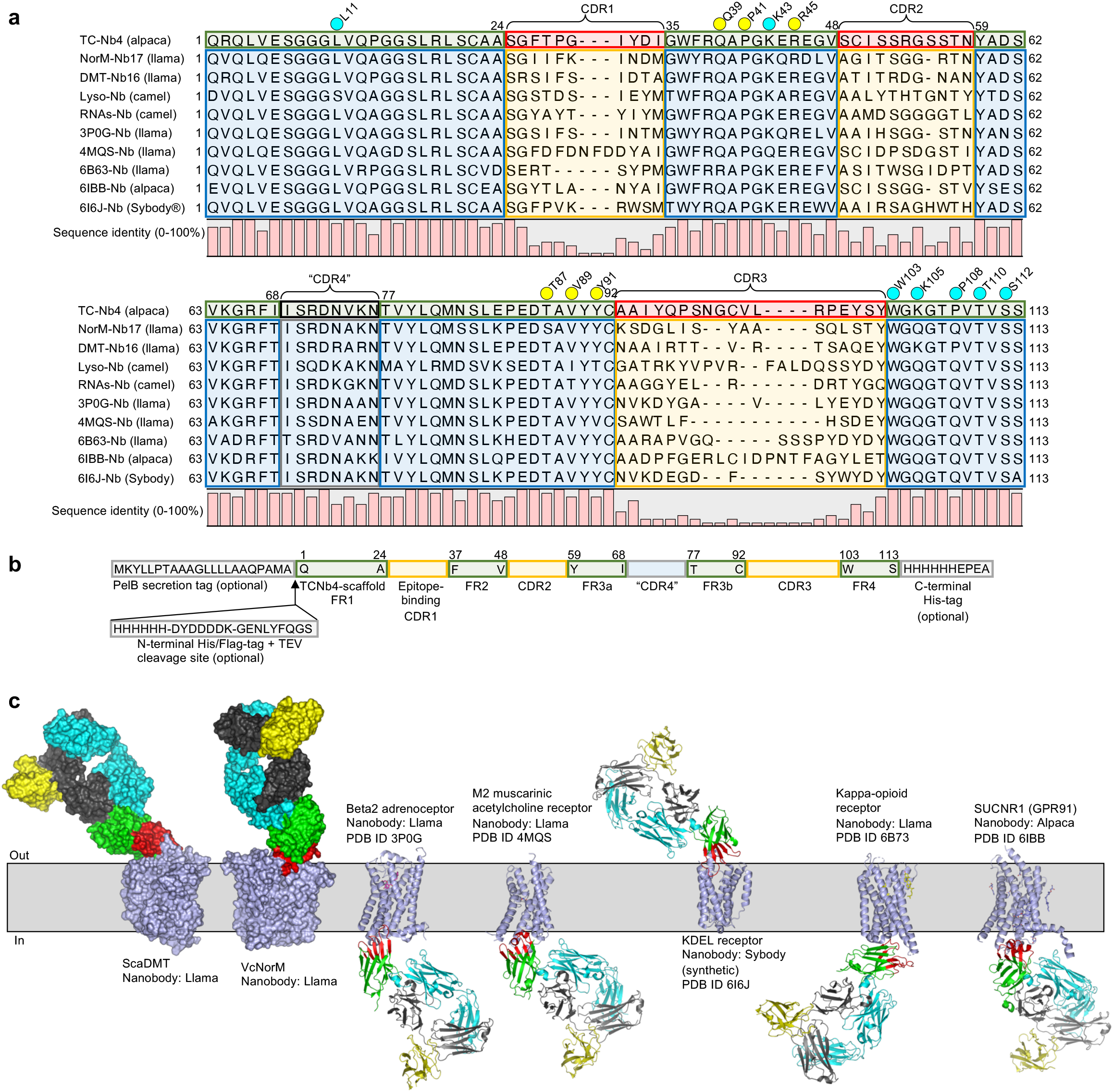
*in silico* CDR grafting and NabFab binding analysis for potential structural targets. **a**, Protein sequence alignment of TC-Nb4(32), NorM-Nb17, DMT-Nb16(37), Lyso-Nb(35), as well as selected Nbs of known membrane protein-Nb complexes(10–12, 49, 50) (PDB ID: 3P0G, 4MQS, 6B73, 6IBB, 6I6J), generated with Clustal Omega(51), in Kabat numbering(47). Green boxes indicate scaffold regions of TC-Nb4. Blue boxes indicate equivalent scaffold regions other nanobodies. Red boxes indicate CDRs of TC-Nb4. Yellow boxes indicated equivalent CDR regions of other Nbs. A black box indicates “CDR4” of TC-Nb4. A grey box indicates the equivalent region in other nanobodies. Note that "CDR4” is a non-variable loop that in some cases interacts with the target protein of the respective Nb. Residues at the Nb-binding interface to NabFab are indicated with colored dots on top. Yellow dots indicate a strong contribution to NabFab binding. Cyan-colored dots indicate a weak contribution, as described in Figure 3b. **b**, Schematic of CDR-grafting strategy: Green boxes indicate regions of the TC-Nb4 scaffold that were retained during grafting. Yellow boxes indicate epitope binding CDR-regions of different Ns that were grafted into the TC-Nb4 scaffold. A grey/blue box indicates “CDR4” which can also be grafted. Flanking grey boxes indicate the secretion and purification tags. Note that the His-tags are not necessary for NabFab-binding. **c**, Structures of complexes of VcNorM and ScaDMT in surface representation, as well as *in silico* docked (PyMol(52)), putative complexes of selected membrane protein-nanobody complexes in ribbon representation. Membrane proteins are colored in blue, original Nb CDRs in red, TC-Nb4-scaffold in green, NabFab HC in cyan, NabFab LC in grey, and anti-Fab Nb in yellow.

### Structural considerations for a universal nanobody Fab

As part of the phage display selection process, we employed an epitope masking strategy to focus the available binding epitopes on the opposing face of the nanobody relative to the CDR loops. However, while the epitope position is important, it is the orientation of the Fab extending from the epitope that is of most consequence. The relative angles defined by the orientation of the Fab to the Nb and the Nb to its membrane target combine to delineate the potential universality of the Fab. In this regard, the NabFab engages the TC-Nb4 scaffold at an obtuse angle with respect to the centroid of the CDR residues, orienting the Fab away from the antigen target. The second factor is the angle of engagement of the Nb to its target. In the examples described here, the chimeric Nbs for VcNorM and ScaDMT bind to their respective targets at significantly different angles, yet the bound NabFab pointed away from the detergent micelle and proved to be an effective fiducial in both cases (Fig. 2). To further explore the potential applicability of NabFab to study other membrane protein-Nb complexes, we performed sequence alignments and structural superpositions with published structures of five different membrane proteins (Fig. 5). The structure-based sequence alignments (Fig. 5a) establish the precise boundaries within the Nb scaffold that define the CDR grafting strategy. Based on these alignments, we propose a generally applicable grafting protocol for nanobodies available for other target proteins (Fig. 5b), ensuring that the resulting chimera will bind NabFab with full affinity. In addition to the hypervariable CDR loops 1-3, we advise to also graft the fourth, invariable loop referred to as “CDR4”. Although this loop is generally conserved, it occasionally contributes to binding. Notably, as neither of the Nb termini is involved in NabFab binding, the location of a purification tag of the chimeric Nb construct can be chosen freely. Finally, we superimposed the Nb component of our VcNorM-containing complex with those of published membrane protein-Nb structures following an *in silico* grafting of the respective Nbs (Fig. 5c). The superpositions suggest that NabFab is neither expected to clash with the target proteins nor with the lipid bilayer. This suggests that NabFab will likely be applicable for the cryo-EM analysis of a wide range of membrane protein-Nb complexes, including GPCRs.

### Conclusions

We have developed and validated a universal cryo-EM fiducial for the study of protein-nanobody complexes. It is a robust and reliable tool based on its ability to strongly and specifically bind the engineered scaffold portion of folded nanobodies. We provide an easy-to-follow grafting strategy that facilitates the repurposing of Nb scaffolds to ensure binding to the universal Fab (NabFab), irrespective of the origin of the Nb. NabFab is conveniently expressed in in *E. coli,* and its synthetic nature allows it to be further engineered to introduce other labels that could be useful for complementary cell biology studies or to facilitate other binding proteins. Our results suggest that NabFab is not only useful for high-resolution structure determination of small (membrane) proteins by cryo-EM, but its shape characteristics with the attached anti-Fab elbow Nb can be powerfully exploited for the rapid assessment of test datasets.

Despite recent advances in the prediction of 3D structures by artificial intelligence-based algorithms(45, 46), determining experimental high-resolution structures of proteins in distinct conformational states and bound to ligands and inhibitors remains essential for basic science and drug development. Many researchers have generated nanobodies against membrane proteins, often trapping specific functional states. NabFab offers a convenient way for those projects to transition from X-ray to cryo-EM or to increase the resolution of cryo-EM studies. NabFab binding also appears compatible with the simultaneous application megabodies(28), which might help to improve particle solubility and orientational distribution. Finally, NabFab might be useful for other applications: The rigid linkage of NabFab to target nanobodies might improve the precision of single-molecule imaging studies using FRET. Moreover, the humanized scaffold of NabFab combined with the low immunogenicity of nanobodies(6) also offers opportunities in medical imaging or drug development.

## Methods

### Expression and purification of nanobodies

TC-Nb4, NorM-Nb17_4, DMT-Nb17_4, PglB-Nb17_4, and TC-Nb11_4 were purified as described previously(32).

BtuF-Nb9_4, Lyso-Nb_4, and RNAs-Nb_4 were cloned in pET26b+ vector as N-terminal Histag fusion constructs. These were expressed in *E. coli* C43(DE3) cells in Terrific Broth media supplemented with 0.4% (v/v) glycerol. The cells were grown until mid-log phase at 37°C, induced with 1 mM IPTG and grown for 16-20 hrs at 25°C for expression of the nanobodies. The overexpressed nanobodies were purified by Ni-NTA chromatography followed by SEC on Superdex75 increase columns. For crystallization trials with the Lyso-Nb_4, the Histag was cleaved with TEV protease followed by a subtractive Ni-NTA chromatography before SEC experiments.

### Enzymatic biotinylation of nanobodies

Nanobodies were biotinylated via C-terminal insertion of the amino acid sequence (GGGS)3-GLNDIFEAQKIEWHE-GGGS-H6. And subsequent BirA mediated biotinylation, as described previously(33). The extent of labeling was verified by streptavidin (SA) pull-down assay.

### Grafting of CDRs into nanobody TCNb4 scaffold

Based on structural comparisons and sequence alignments in Clustal Omega(51) of different nanobodies we decided in a grafting strategy as outlined in Fig. 4 b. For expression in *E. coli* cells all nanobodies contained an N-terminal PelB secretion tag (MKYLLPTAAAGLLLLAAQPAMA-Nanobody), which is naturally cleaved during bacterial expression. According to Kabat numbering(47) we combined residues 1-24, 35-48, 59-68, 77-92, and 103-113 of TC-Nb4, with the respective CDRs of the nanobody of interest. For NorM-Nb17_4 “CDR4” of TC-Nb4 was retained as it was very similar to NorM-Nb17’s “CDR4”. For purification we either (i) inserted an N-terminal His6-Flag-TEV cleavage site (PelB-HHHHHH-SSDYKDDDDK-GENLYFQGS-Nanobody) or we used (ii) a C-terminal His6-CaptureSelect^TM^ C-tagXL (PelB-Nanobody-HHHHHH-EPEA). (i) was used for ButF-Nb9_4, Lyso-Nb_4, and RNAs-Nb9_4, (ii) for TC-Nb4, NorM-Nb17_4, DMT-Nb17_4, and PglB-Nb17_4.

### Phage display selection

Selection for TC-Nb4 was performed as described. In the first round of selection, 200 nM of target was immobilized on 250 μl streptavidin (SA) magnetic beads. This was followed by rigorous washing steps to remove unbound protein followed by a 5 min incubation with 5 μM D-biotin to block unoccupied SA sites on the beads to prevent nonspecific binding of the phage pool. The beads were then incubated with the phage library (Miller et al., 2012), containing 10^12^–10^13^ virions/ml for 30 min at RT with gentle shaking. The resuspended beads containing bound phages after extensive washing were used to infect freshly grown log phase *E. coli* XL1-Blue cells. Phages were amplified overnight in 2YT media with 50 μg/mL ampicillin and 10^9^ p.f.u./mL of M13 KO7 helper phage. To increase the stringency of selection, three additional rounds of sorting were performed with decreasing target concentration in each round (2^nd^ round: 100 nM, 3^rd^ round: 50 nM and 4^th^ round: 10 nM) using the amplified pool of virions of the preceding round as the input. Unlike solid capture in the first round, the 2^nd^ – 4^th^ rounds involve solution capture. These rounds are carried out in a semi-automated platform using the Kingfisher instrument. 2^nd^ round onwards, 5 uM of TC:CD320 were used in all the binding and washing steps to prevent the phage pool to bind to the CDRs of TC-Nb4. From 2^nd^ round, the bound phages were eluted using 0.1 M glycine; pH 2.7. This technique often risks the enrichment of nonspecific and SA binders. In order to eliminate them, the precipitated phage pool were negatively selected against 100 μL of SA beads before each round. The “precleared” phage pool was then used as an input for next round in the campaign.

### Enzyme-linked immunoabsorbent assays (ELISA)

All ELISA experiments were performed in a 96-well plate coated with 50 μL of 2 μg/mL neutravidin in Na2CO3; pH 9.6 and subsequently blocked by 0.5% in PBS. A single-point phage ELISA was used to rapidly screen the binding of the obtained clones in phage format. Colonies of *E. coli* XL1-Blue harboring phagemids were inoculated directly into 500 μL of 2YT broth supplemented with 100 μg/ml ampicillin and M13 KO7 helper phage. The cultures were grown at 37 °C for 16-20h in a 96-deep-well block plate. Culture supernatants containing the Fab phage were diluted 20-fold in ELISA buffer and transferred to ELISA plates that were incubated with 50 nM of biotinylated TC-Nb4 in experimental wells and buffer in control wells for 15 min at RT. The ELISA plates were incubated with the phage for another 15 min and then washed with PBST. The washed ELISA plates were incubated with HRP-conjugated anti-M13 mouse monoclonal antibody (1:5000 dilution in PBST) for 30 min. The plates were again washed, developed with TMB substrate and quenched with 1.0 M HCl, and absorbance (A_450_) was determined. The background binding of the phage was monitored by the absorbance from the control wells.

Protein based single point ELISA was performed to determine the cross-reactivity of the Fabs to the different nanobodies in the grafted TC-Nb4 scaffold. 50 nM of TC-Nb4 and BtuF-Nb9_4, Lyso-Nb_4 and RNAs-Nb_4 were immobilized on ELISA plate followed by incubation wit 250 nM of the purified Fabs 1, 2, 6, 12, 14, 16, 18, 19, 21, 23, 28. The plates were washed and the bound antigen-sAB complexes were incubated with a secondary HRP-conjugated protein L (1:5000 dilution in PBST). As with phage ELISA, the plates were again washed, developed with TMB substrate and quenched with 1.0 M HCl, and absorbance (A_450_) was determined.

### Cloning, overexpression and purification of Fabs

*E. coli* C43 cells were transformed with Fabs cloned in the expression vector pRH2.2. Fabs were grown in TB autoinduction media with 100 μg/mL ampicillin at 25°C for 16 hrs. Harvested pellets were resuspended in PBS, supplemented with 1 mM PMSF, 1 μg/mL DNase I. The suspension was lysed by ultrasonication. The cell lysate was incubated at 65°C for 30 min. Heat-treated lysate was then cleared by centrifugation, filtered through 0.22 μm filter and loaded onto a 5 ml HiTrap protein L column pre-equilibrated with 20 mM Tris; pH 7.5, 500 mM NaCl. The column was washed extensively with 20 mM Tris; pH 7.5, 500 mM NaCl followed by elution of Fabs with 0.1 M acetic acid. The eluted protein was directly loaded onto a 1 ml ResourceS column pre-equilibrated with 50 mM NaOAc; pH 5.0. The column was washed with the equilibration buffer and Fabs were eluted with a linear gradient 0–50% of 50 mM NaOAc; pH 5.0, 2 M NaCl. Purified Fabs were dialyzed overnight against 20 mM HEPES; pH 7.5, 150 mM NaCl. The quality of purified Fabs was analyzed by SDS-PAGE.

### Binding kinetics by SPR

All SPR experiments were performed at 20°C using MASS-1 (Bruker) instrument. 20 nM TC-Nb4 and BtuF-Nb9_4, Lyso-Nb_4, and RNAs-Nb_4 were immobilized onto a nitrilotriacetic acid (NTA) sensor chip via His-tag. 2-fold serial dilutions of Fab19 and Fab14 were injected following ligand immobilization on the sensor chip. For each kinetic experiment at least five dilutions of the Fabs were tested. The kinetic parameters were determined by fitting the data in 1:1 Langmuir model.

### Expression and purification of ScaDMT

ScaDMT was expressed as described previously(37) with minor modifications. All steps of the purification were performed at 4 °C or on ice. Cells were resuspended in 25 mMEPES-NaOH pH 7.4, 150 mM NaCl, 0.1 mg/ml DNAseI and disrupted in a M-110-l microfluidizer (Microfluidics) at 15,000 p.s.i. chamber pressure. Cell debris were pelleted by centrifugation at 10,000 x g for 20 min. The supernatant was decanted, and membranes were pelleted by centrifugation at 100,000 x g for 1h. The membranes were resuspended in 25 mM HEPES-NaOH pH 7.4, 150 mM NaCl 10% glycerol, 1% DDM at a concentration of 3 ml per gram cell pellet used. After incubation for 30 min remaining debris were pelleted by centrifugation at 30,000 x g for 30 min and ScaDMT was purified by affinity chromatography (IMAC). Finally, the buffer was exchanged via a PD-10 desalting column (GE Healthcare) into 10 mM HEPES-NaOH pH 7.4, 150 mM NaCl, 3% glycerol, 0.015% DDM.

### Expression and purification of VcNorM

All media were supplemented with 100 μg / ml ampicillin and 1 % (w/v) glucose. E. coli BL21 DE3 gold cells harboring the expression plasmid were grown on LB agar plates for ~10 hours at 37 °C. A single colony was picked with a sterile toothpick that was dropped into 500 ml room temperature TB medium in a baffled 2-liter Erlenmeyer flask. The preculture was grown for 10 to 12 hours at 37 °C and 80 rpm. At an OD600 of 1, 50 ml of the preculture were transferred to baffled 5-liter Erlenmeyer flasks containing 1.5 liters of pre-warmed TB medium. Cells were grown at 37 °C and 120 rpm and expression of the protein was induced with 0.2 mM IPTG at an OD600 of 3.0 After 60 minutes, cells were harvested by centrifugation at 15’000 x g for 10 minutes at 4 °C.

All steps were carried out at 4 °C unless stated differently. Cells were resuspended in 50 mM Tris-HCl pH 7.5, 200 mM NaCl at a ratio of 1 : 5 (g cells : ml buffer). Cells were cracked by three passes through a 200 μm chamber of an M-110L microfluidizer (Microfluidics) at 15,000 psi external pressure. The lysate was cleared by centrifugation at 4’000 x g for 20 minutes and pelleted by ultracentrifugation at 142,000 x g for 35 minutes. Membranes were resuspended in 50 mM Tris-HCl pH 7.5, 200 mM NaCl, 20 mM imidazole-HCl pH 8.0 at a ratio of 1 : 2 (g cells : ml final volume). DDM was added for solubilization to a final concentration of 1 % (w/v). After stirring the suspension for 90 minutes, insoluble material was removed by centrifugation for 30 minutes at 40’000 x g and the supernatant decanted. VcNorM was purified by metal affinity chromatography (IMAC). The buffer was exchanged to 10 mM Tris-HCl pH 7.5, 100 mM NaCl, 0.5 mM EDTA-NaOH pH 8.0, 0.016 % (w/v) DDM or to the buffer specified in the results section using a PD-10 desalting column (GE-Healthcare)

### Nanobody generation against VcNorM

VcNorM was incorporated into liposomes consisting of a 3 : 1 (w/w) mixture of E. coli polar lipid extract and L-α-phosphatidylcholine as described previously(53). A llama (Llama glama) was injected 6 times with liposome reconstituted VcNorM. Peripheral blood lymphocytes were isolated from samples collected 3 to 4 days after the last immunization and nanobodies were generated as described previously(40).

### Complex formation and EM sample preparation

All steps of were performed at 4 °C or on ice. Complex formation for VcNorM was achieved by mixing 14 μM VcNorM, 7 μM NorM-Nb17_4, 7.7 μM NabFab, and 11.2 μM Anti-Fab Nb and subsequent incubation overnight with gentle agitation. Comblex formation for ScaDMT was achieved by mixing 4.5 μM ScaDMT, 4.95 μM DMTNb16_4, 5.4 μM NabFab, and 6.75 μM Anti-Fab nanobody.

The complexes were concentrated using a centrifugal filter (100 kDa cutoff, Amicon) and loaded on a Superdex S200 increase column using the respective desalting buffer as a running buffer but at 0.08% DDM. Fractions containing the respective membrane proteins were collected for Cryo-EM grid preparation.

The final VcNorM complex sample was at a concentration of 7.1 mg/ml and was supplemented with 0.006% cholesteryl hemi-succinate (CHS). The final ScaDMT complex sample was at a concentration of 6.4 mg/ml and was not further supplemented. Quantifoil holey carbon grids, Cu, R 1.2/1.3, 300 mesh, were glow discharged for 45 s, 25 mA using a PELCO easiGLOW glow discharger. Sample (2.0 μl) was applied to the cryo-EM grids and blotted for 2.5–3.5 s before plunge freezing in a liquid ethane–propane mixture with a Vitrobot Mark IV (Thermo Fisher Scientific) operated at 4 °C and 100% humidity.

### EM data collection

Data was recorded on a Titan Krios electron microscope (Thermo Fischer Scientific) operated at 300 kV, equipped with a Gatan BioQuantum 1967 filter with a slit width of 20 eV and a Gatan K3™ camera. Movies were collected semi-automatically using SerialEM(54) at a nominal magnification of 130,000 and a pixel size of 0.335 Å (VcNorM) and 0.33 Å (ScaDMT) per pixel in super-resolution mode. The defocus range was −0.6 to −2.8 μm. Each movie contained 40 images per stack with a dose per frame of 1.3 electrons/Å^2^.

### EM data processing, model building, and refinement

For the test structure of the VcNorM complex, first a test dataset of 380 movies was collected and corrected for beam induced motion correctio using MotionCor2(55). 139,608 particles were auto-picked in Laplacian of gaussian mode (LOG) and extracted with threefold binning at 1.98 Å/pixel. After 2 rounds of 2D classification 10,193 particles were clearly attributed to the protein complex of interest and an initial model was calculated thereof. After one subsequent round of 3D classification 8,514 particles were selected and refined to 12.6 Å resolution.

For the high-resolution structure of the VcNorM complex 13,009 movies were collected and corrected for beam induced motion correctio using MotionCor2(55). 9,711 micrographs were selected for further processing in RELION 3.1(42). The contrast transfer function was estimated using Gctf(56). In a combined approach of LOG-based and 2D class reference-based particle picking a total of 6,365,855 particles were auto-picked and were sorted by 2D and 3D classification. 483,559 particles were re-extracted to 0.66 Å/pixel and were subjected to another round of 3D classification. Therefrom, 433,559 particles were selected and subjected to particle polishing and per-particle CTF refinement. The particles were refined to 3.68 Å resolution. Moving the fulcrum from the TMDs of VcNorM into the chimeric nanobody improved the overall resolution to 3.47 Å. Local resolution resolution estimates were generated in RELION 3.1(42).

For the test structure of the ScaDMT complex, first a test dataset of 625 micrographs was collected and corrected for beam induced motion correctio using MotionCor2(55). 106,551 particles were auto-picked (LOG) and extracted with threefold binning at 1.98 Å/pixel. After one round of 2D classification a class containing 3,659 particles was clearly attributed to the protein complex of interest and an initial model was calculated thereof. After one subsequent round of 3D classification with 20,494 particles included, 10,851 particles were selected and refined to 12.3 Å resolution.

For the high-resolution structure of the ScaDMT complex 21,251 movies were collected and corrected for beam induced motion correctio using MotionCor2(55). 17,957 micrographs were selected for further processing in RELION 3.1(42). The contrast transfer function was estimated using Gctf(56). 2,416,345 particles were auto-picked (LOG) and were extracted at 2.64 Å/pixel. After sorting of the particles by 2D and 3D classification, 404,841 particles were re-extracted at 0.66 Å/pixel and were subjected to particle polishing and per-particle CTF refinement. The particles were refined to 3.99 Å resolution. After one round of heterogeneous refinement in cryoSPARC v3.2(48) 314,541 particles were selected and refined to 3.78 Å resolution by non-uniform refinement(57) with optimized per-particle defocus and optimized CTF per-group parameters. For comparability, local resolution estimates were calculated in RELION 3.1(42).

The structure of the dimeric complex of ScaDMT was determined based of a subclass of 40,512 particles of that appeared during 3D classification of the ScaDMT complex. After one subsequent round of 3D classification 34,100 particles were selected refined to 8.45 Å resolution.

Model building was performed in Coot(58) and models were refined in PHENIX(59). The structure of VcNorM was built de-novo. The structures of the other components were built based on rigid body docking of the crystal structures of TCNb4(32), and the Fab-anti-Fab nanobody complex(31). The structure of the ScaDMT:nanobody complex was built based on the published crystal structure thereof (PDB ID 4WGV)(37), and the Fab-anti-Fab nanobody complex(31). Due to lower map quality in the respective regions, the part of the crystal structure containing the constant part of the Fab and the anti-Fab nanobody was rigid body docked into the map after the last round of refinement. For the structure of the homo-dimeric ScaDMT complex, the structure of the monomeric complex was docked into the map in UCSF Chimera(60) and was not further refined.

### Crystallization, data collection, structure determination and refinement statistics for Lyso-Nb_4:NabFab complex

NabFab was incubated with 1.5 molar excess of Lyso-Nb_4 on ice for 30 mins and the complex was subjected to SEC on an S75 increase column whereby the 1:1complex of the Fab-Nb complex was separated from the excess Nb. Monodispered peak fractions of the complex in 10 mM HEPES, 100 mM Sodium chloride, pH 7.5 after SEC were concentrated to 10 mg/mL and subjected to crystallization trials by hanging-drop vapor diffusion technique using robotic system Mosquito crystal (SPT Labtech Inc) and commercially available crystallization screens including PACT, PEGs, Protein Complex Suite (NeXtal) and PEG/Ion Suite (Hampton Research). Precipitant solution from crystallization screen was dispensed at 1:1 macromolecular complex/precipitant solution ratio in a total drop volume of 400 nL at room temperature. Crystals of NabFab:Lyso-Nb_4 were obtained in conditions containing 0.02 M Citric acid, 0.08 M BIS-TRIS propane, pH 8.8, 16 % (w/v) PEG3350 (PEG/Ion Suite - H5). Single crystals of Fab-antiLysNb complex were soaked in cryoprotectant solution containing 0.2 M Ammonium iodide, 20 % (w/v) PEG3350, and 10 % (w/v) Glycerol. Prior to data collection crystals were flash-frozen and stored in liquid nitrogen.

Two datasets were collected remotely from a single crystal of NabFab:Lyso-Nb_4 at the Northeastern Collaborative Access Team (NE-CAT) on the 24-ID-E beamline at Argonne National Laboratory (Argonne, IL) at 100 K. Individual datasets were indexed, integrated in *XDS(61, 62)* and scaled in *Aimless(63)* using RAPD internet based platform containing a modular package of programs at NE-CAT website. Two datasets were merged and used for phasing by molecular replacement method in the program *Phaser(64)*. Crystal structure of nanobody D3-L11 (PDB ID 6JB9) and homologous antibody fragment (PDB ID 5C8J) were used as starting templates for molecular replacement. Structure was refined by *phenix.refine(65)* and manually corrected in *Coot(58, 66)* The coordinates and structure factors were uploaded into the Protein Data Bank as entry 7RTH. The data collection, structure determination and refinement statistics are summarized in the Supplementary Data Table 4.

### Figure preparation and data analysis

Figures of structural representations were prepared in PyMol(52), UCSF Chimera(60), and UCSF ChimeraX(67). Graphs were prepared in GraphPad Prism 9. Protein sequence alignments were generated by Clustal Omega(51) and were depicted using CLC Genomics Workbench 12.

## Acknowledgments

This research was supported by the Swiss National Science Foundation (SNF) grant 310030_189111 to K.P.L., and National Institutes of Health grant GM117372 to A.A.K. Cryo-EM data was collected at the ScopeM facility at ETH Zürich, and we thank the staff of ScopeM for technical support. We thank R. Dutzler for providing the expression plasmid and purification protocols for ScaDMT, M. Mikolin for technical support with protein expression and purification, and R.N. Irobalieva for help with EM data collection. The x-ray diffraction data were collected at NECAT beamline sector 24-ID-E (P30 GM124165), with the Eiger 16M detector (S10OD021527) at the Advanced Photon Source (DE-AC02-06CH11357).

## Author contributions

J.S.B. and K.P.L. conceived the project. J.S.B., S.M., A.A.K. and K.P.L. designed experiments. S.M. performed phage-display selection, generated, expressed and purified NabFab, and performed ELISA, SPR and aSEC-based characterization. A.A.K. provided the Chaperone-Assisted Structure Determination phage display pipeline and supervised synthetic antibody generation. J.S.B. generated biotinylated nanobodies, designed grafted nanobody constructs, generated EM samples, calculated EM structures, and performed model building and refinement. J.K. collected EM data. M.N. cloned VcNorM construct, established original purification protocol, and generated VcNorM sample for nanobody production. E.P. generated NorM-Nb17 under supervision of J.S., E.V.F. and S.M. determined the crystal structure of the Lyso-Nb_4:NabFab complex and performed model building and refinement thereof. J.S.B., S.M., A.A,K, and K.P.L. analyzed data and wrote the paper.

## Data availability

Atomic coordinates of the complexes ScaDMT:Nb16_DMT-4:NabFab:Anti-Fab Nb and VcNorM:NorM-Nb17_4:NabFab:Anti-Fab Nb models were deposited in the RCSB Protein Data Bank (PDB) under accession number 7PIJ for the ScaDMT monomer, 7PHQ ScaDMT dimer, and 7PHP for VcNorM. The three-dimensional cryo-EM maps were deposited in the Electron Microscopy Data Bank (EMDB) under accession numbers EMD-13438 for ScaDMT monomer, for ScaDMT dimer, and EMD-13425 (Map1, fulcrum in micelle) and EMD-13424 (Map2, fulcrum in NorM-Nb17_4) for NorM. The crystal structure of the Lyso-Nb_4:NabFab complex was deposited in the PDB under accession number 7RTH.

---

**Supplementary Data Fig. 1.**
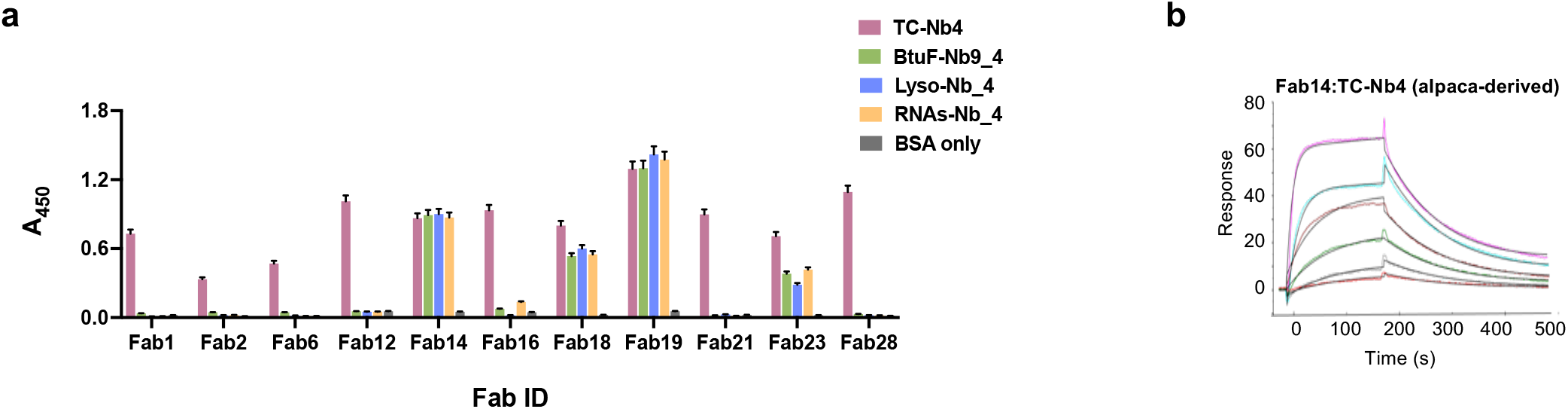
NabFab binding characteristics. **a,** Cross-reactivity of the Fabs tested against TCNb4, BtuF-Nb9_4, Lyso-Nb_4 and RNAs-Nb_4 analyzed by single point protein ELISA. Data are represented as mean ± standard deviation. **b**, SPR sensograms of the binding kinetics of Fab14 with TCNb4.

**Supplementary Data Fig. 2.**
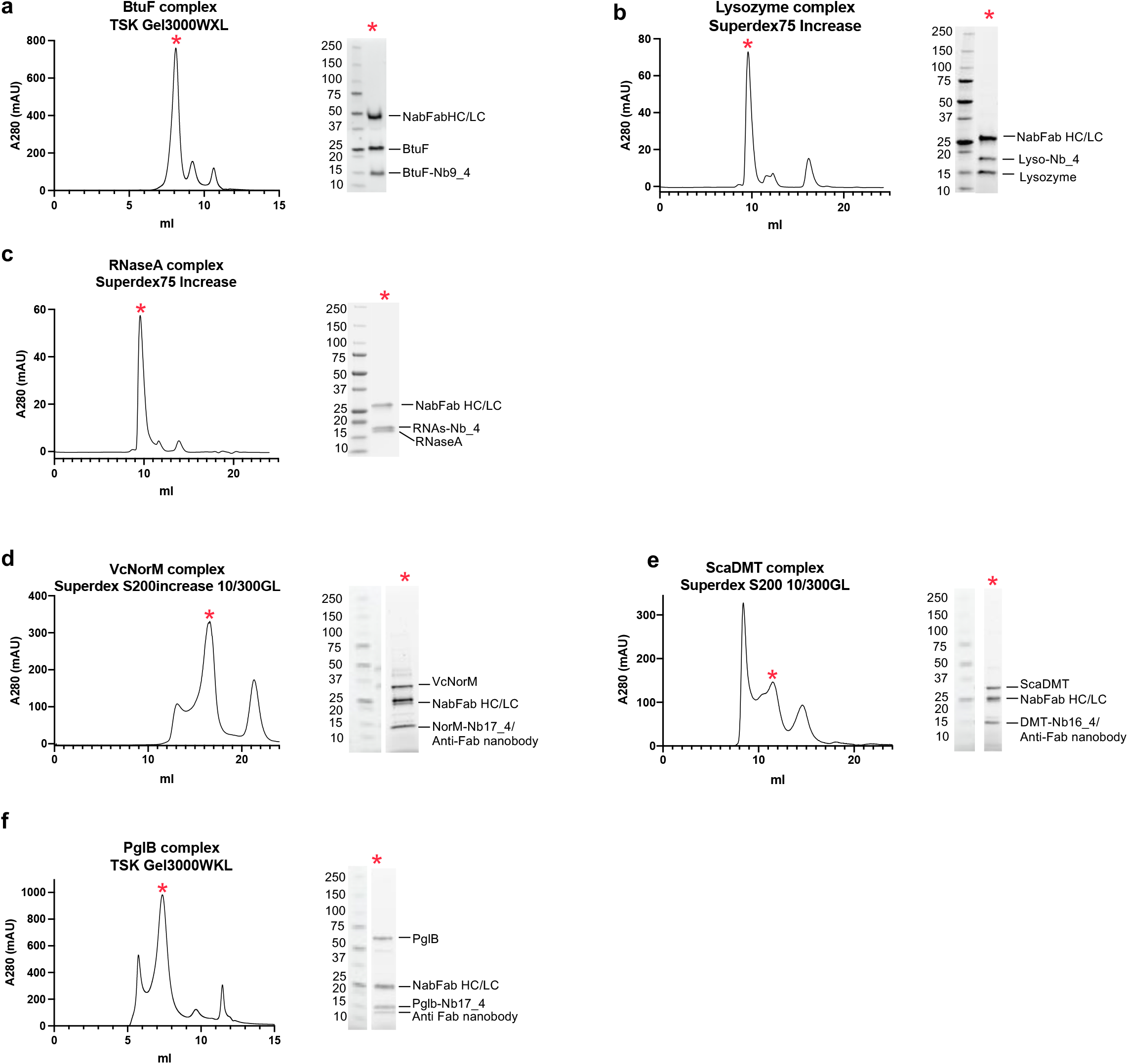
NabFab-nanobody-protein complex formations. SEC profiles and Reducing SDS-PAGE of peak fractions as indicated with a red asterisk fron NabFab chimeric-nanobody complexes with the soluble proteins BtuF (**a**), and lysozyme (**b**), RNAseA (**c**), and with the membrane proteins VcNorM (**d)**, ScaDMT (**e)**, and PglB (**f**).

**Supplementary Data Fig. 3.**
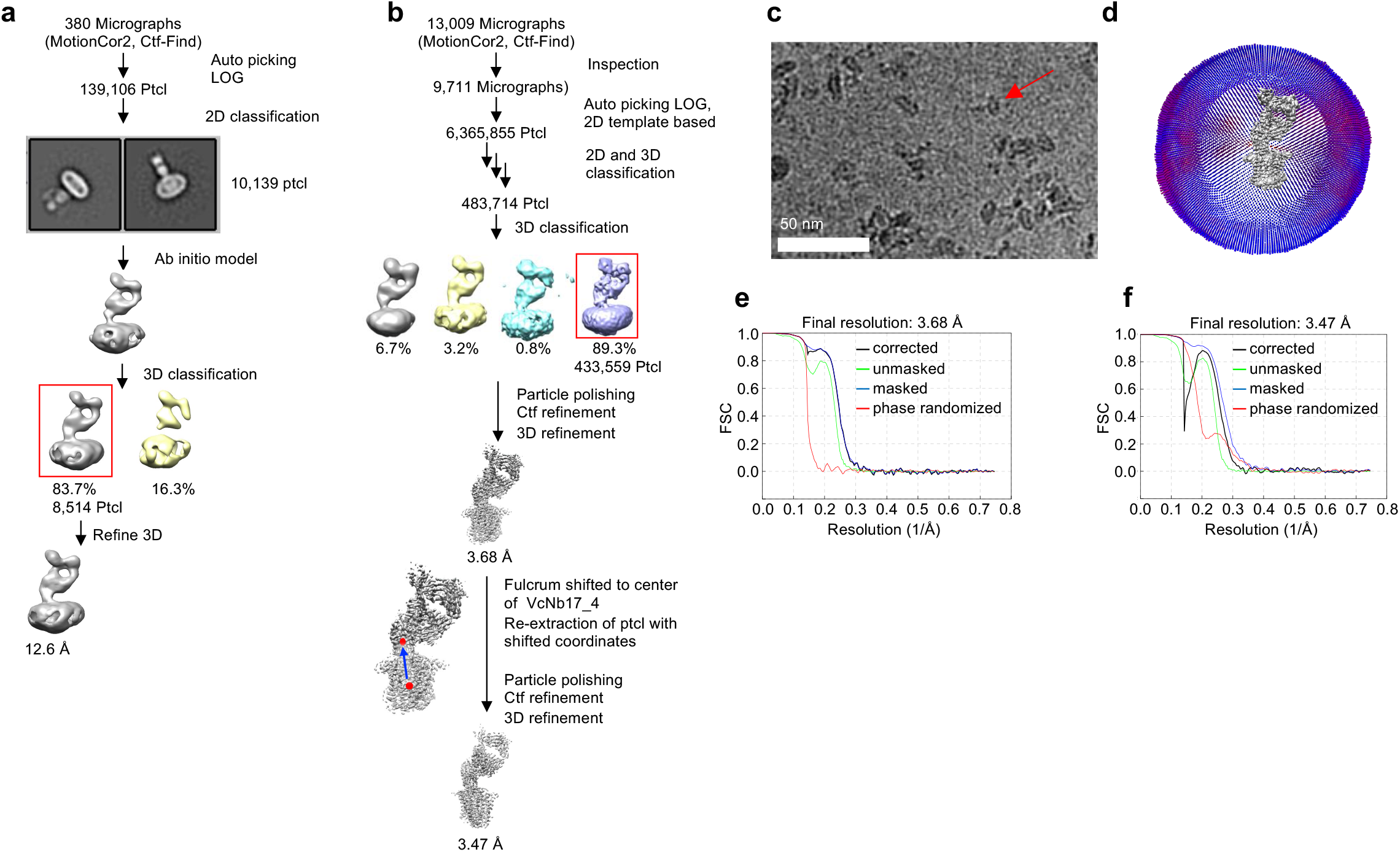
Data processing and structure determination of detergent reconstituted complex of VcNorM, NorM-Nb17_4, NabFab, and Anti-Fab Nb. **a**, Overview of the data processing of a cryoEM test dataset. **b**, Overview of the data processing and structure determination the VcNorM complex. **c**, representative cryoEM micrograph. **d**, Spatial distribution of particles in the final iteration of 3D refinement. **e**, Resolution estimation of the final map via Fourier shell correlation (FSC), with fulcrum positioned in TMD. **f**, Resolution estimation of the final map via Fourier shell correlation (FSC), with fulcrum positioned in NorM-Nb17_4. Unless stated differently, all calculations were performed in RELION 3.1(42).

**Supplementary Data Fig. 4.**
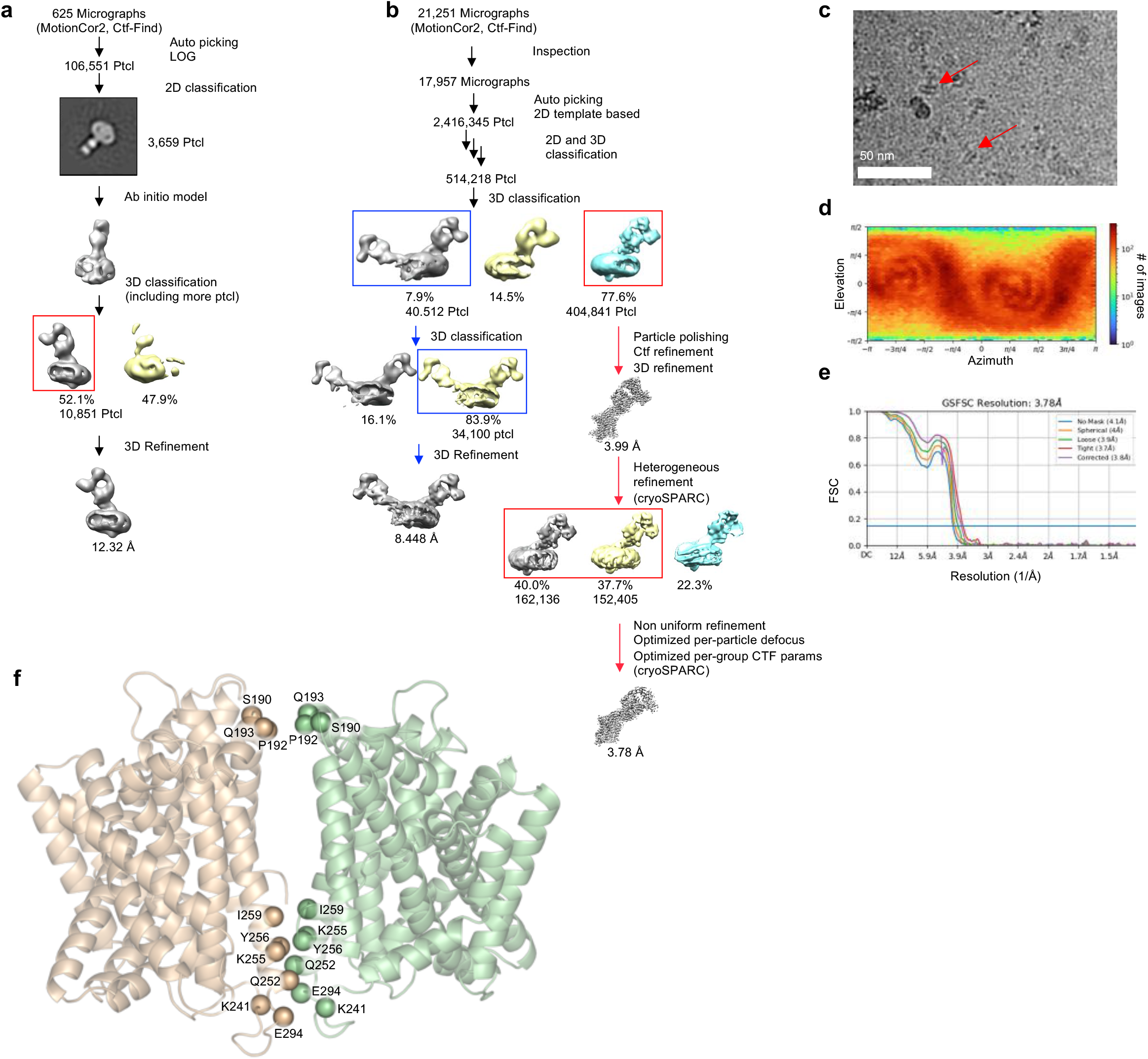
Data processing and structure determination of detergent reconstituted complex of ScaDMT, DMT-Nb16_4, NabFab, and Anti-Fab Nb. **a**, Overview of the data processing of a cryoEM test dataset in RELION 3.1(42). **b**, Overview of the data processing and structure determination the ScaDMT complex and ScaDMT dimer. In RELION 3.1(42) and in cryoSPARC v3.2(48) **c**, representative cryoEM micrograph. **d**, Spatial distribution of particles in the final iteration of 3D refinement as calculated in cryoSPARC v3.2(48). **e**, Resolution estimation of the final map via Fourier shell correlation (FSC), as calculated in cryoSPARC v3.2(48) **f**, Ribbon representation of homo-dimeric structure of ScaDMT with C-alpha positions of residues potentially involved in dimerization shown as spheres.

**Supplementary Data Fig. 5.**
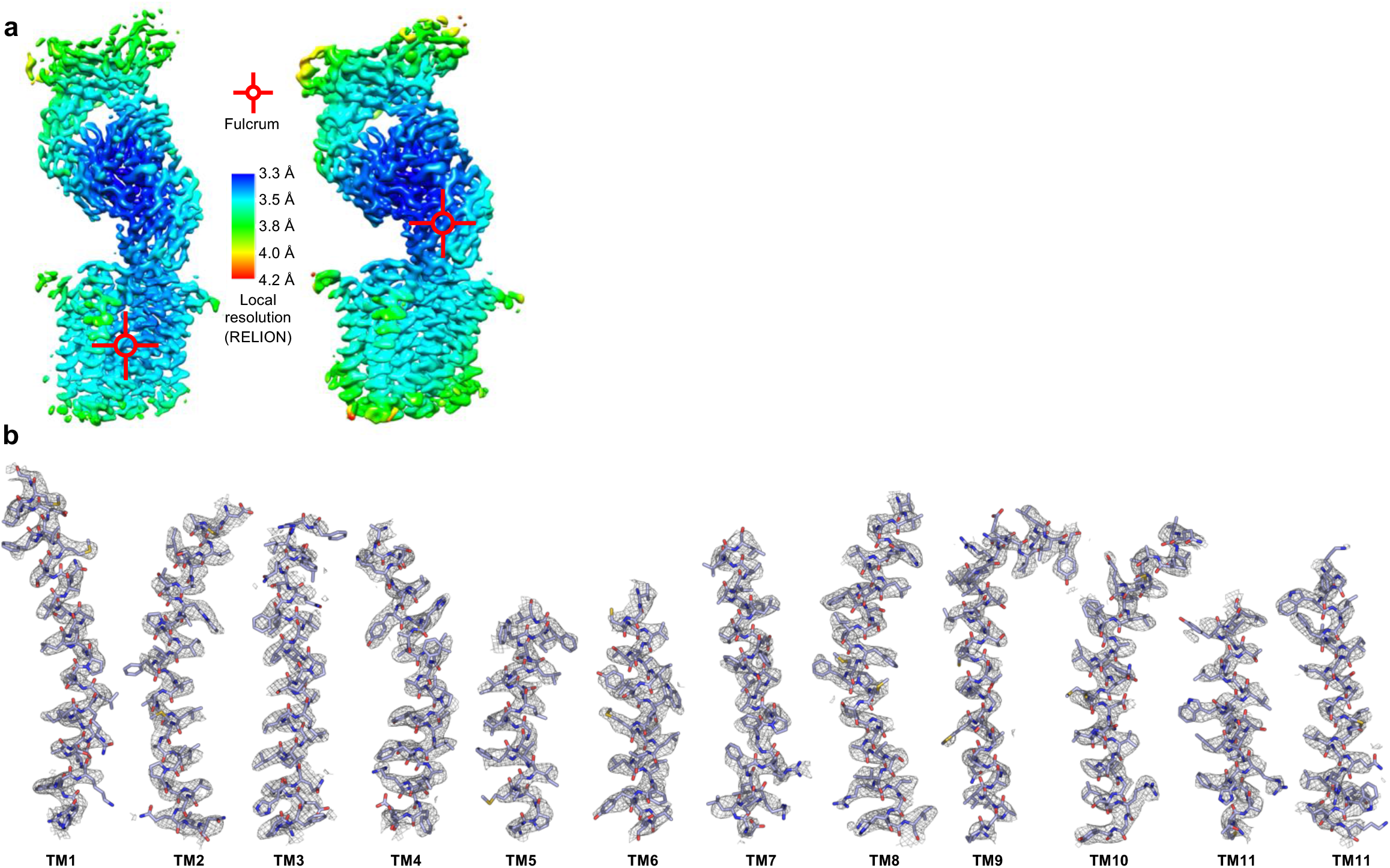
EM map quality assessment of VcNorM complex. **a**, Local resolution estimation of VcNorM maps as calculated in RELION 3.1(42). Red crosshairs indicate the fulcrum during the refinement of the respective map. **b**, EM-density for TM helices of VcNorM for map 1 calculated with fulcrum in micelle (**a**, left). Maps are displayed at a contour level of 10 and were carved at 2.0 Å.

**Supplementary Data Fig. 6.**
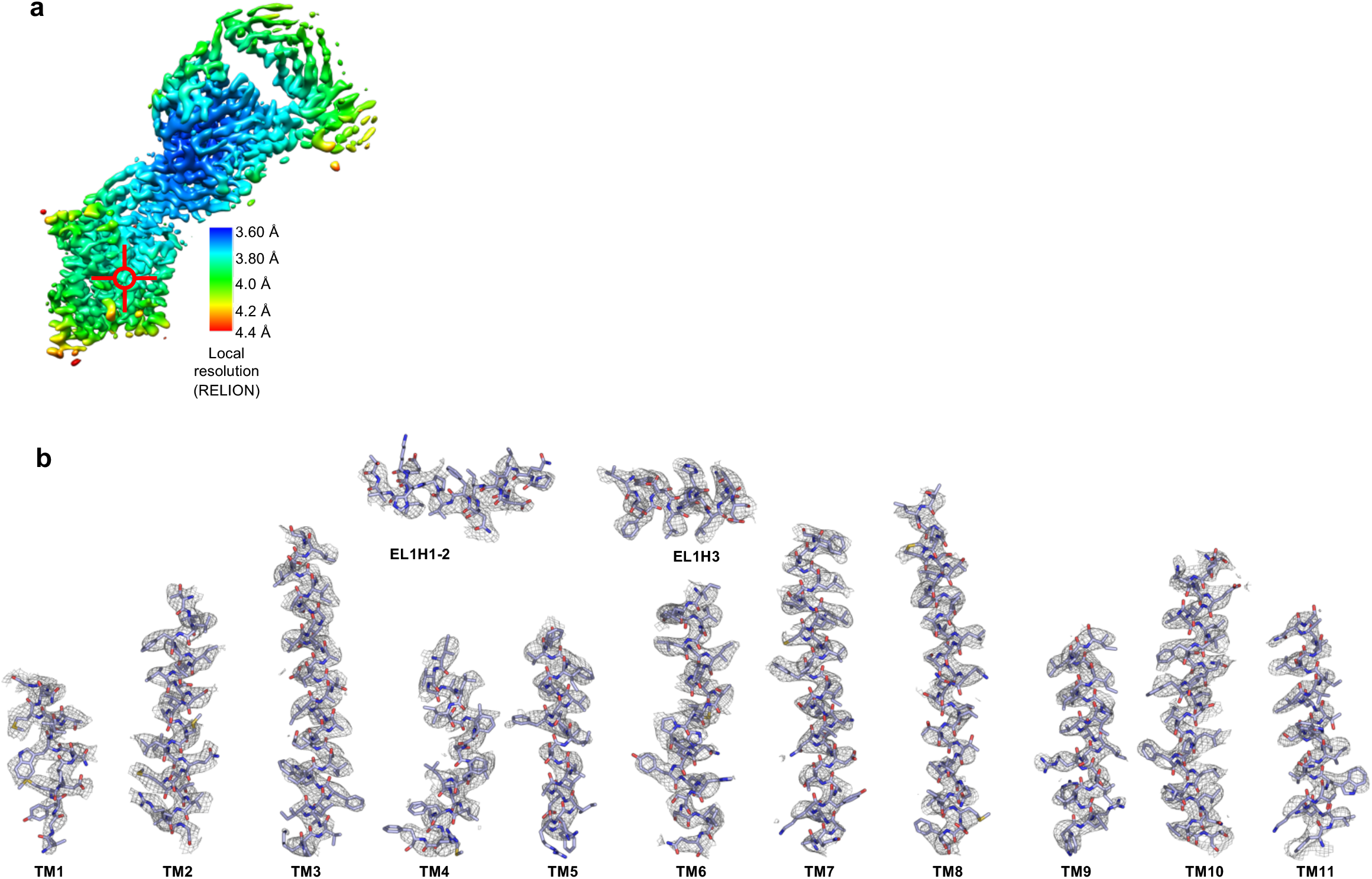
EM map quality assessment of ScaDMT complex. **a**, Local resolution estimation of ScDMT map as calculated in RELION 3.1(42). Red crosshairs indicate the fulcrum during the refinement of the map. **b**, EM-density for TM helices and external helices (ELH) of ScaDMT. Maps are displayed at a contour level of 10 and were carved at 2.0 Å, except for TM4 (rmsd =6) and TM5 (rmsd=8) which required lower rmsd thresholds due to their increased mobility.

**Supplementary Data Fig. 7.**
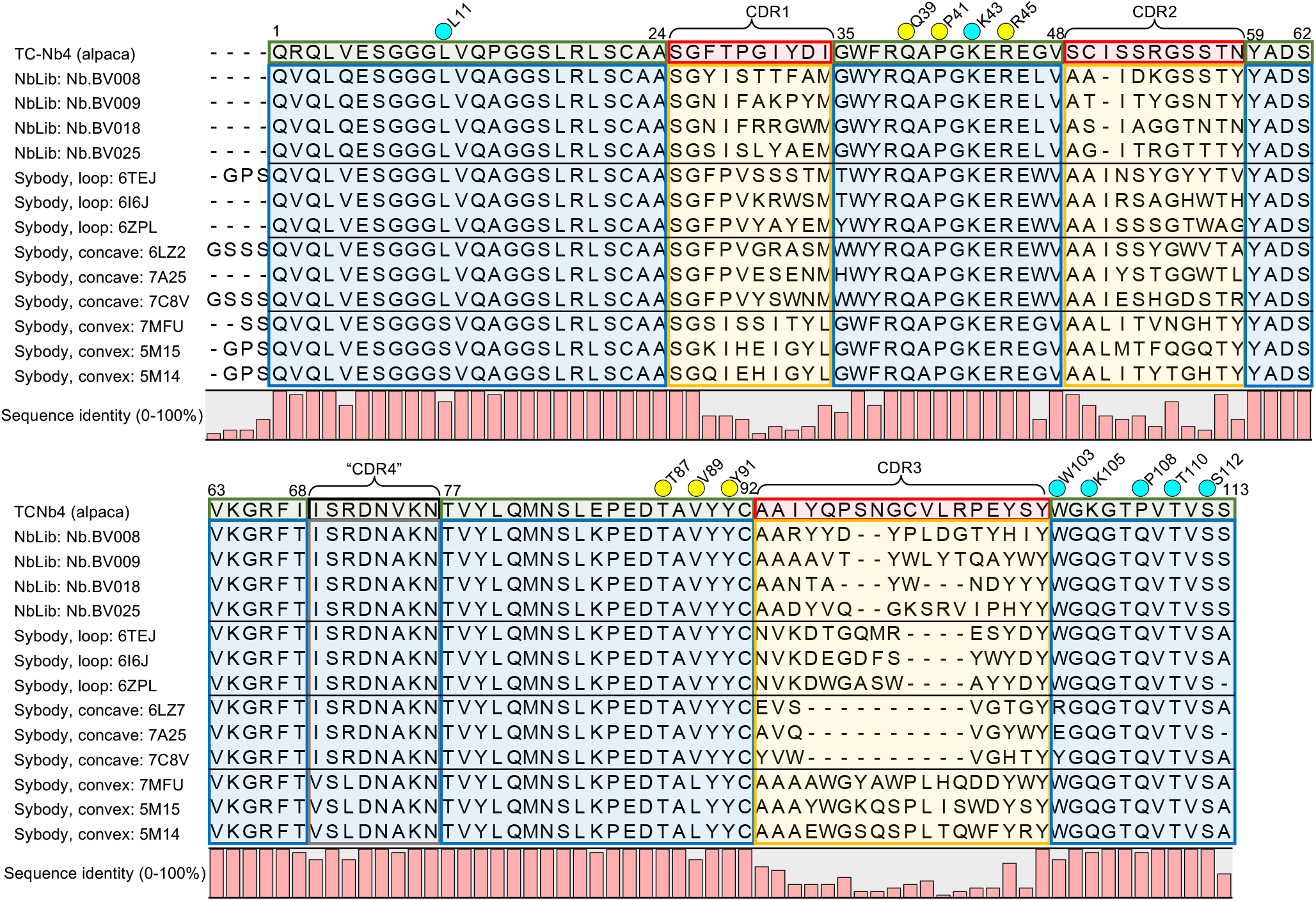
TC-Nb4 scaffold alignments with scaffolds of synthetic Nb libraries. **a**, Protein sequence alignment of TC-Nb4(32), with representative nanobodies from the synthetic nanobody libraries NbLib(43) and Sybody(44). For Sybody scaffolds from all three CDR types were included. The alignments were generated with Clustal Omega(51), in Kabat numbering(47). Labeling and color coding as in Figure 4a.

**Supplementary Data Table 1.** Kinetic and thermodynamic parameters of NabFab binding to wt and Ala mutants of Lyso-Nb_4.

| Nanobody variants | $k_{on}$ ($M^{-1}s^{-1}$) | $k_{off}$ ($s^{-1}$) | $K_D$ (nM) | $KD_{mut}:KD_{wt}$<br>ratio | $\Delta\Delta G^{\circ}_{wt-mut}$<br>(kcalmol$^{-1}$) |
| --- | --- | --- | --- | --- | --- |
| wt | $2.2 \times 10^5$ | $3.5 \times 10^{-4}$ | 1.6 | - | - |
| L11A | $2.0 \times 10^5$ | $8.5 \times 10^{-4}$ | 4.1 | 3 | 0.6 |
| Q39A | $1.4 \times 10^5$ | $3.0 \times 10^{-3}$ | 21.6 | 14 | 1.5 |
| P41A | $8.2 \times 10^4$ | $1.4 \times 10^{-3}$ | 655.1 | 409 | 3.6 |
| K43A | $1.8 \times 10^5$ | $7.4 \times 10^{-4}$ | 4.18 | 3 | 0.6 |
| R45A | $1.6 \times 10^5$ | $6.6 \times 10^{-3}$ | 42.2 | 26 | 1.9 |
| T87A | $1.9 \times 10^5$ | $3.3 \times 10^{-3}$ | 16.4 | 10 | 1.4 |
| V89A | $5.3 \times 10^4$ | $8.9 \times 10^{-3}$ | 1007.1 | 629 | 3.8 |
| Y91A | $1.8 \times 10^5$ | $4.7 \times 10^{-3}$ | 26.9 | 17 | 1.7 |
| W103A | $1.6 \times 10^5$ | $4.9 \times 10^{-4}$ | 3.1 | 2 | 0.4 |
| K105A | $2.1 \times 10^5$ | $1.3 \times 10^{-3}$ | 6.1 | 4 | 0.8 |
| P108A | $1.5 \times 10^5$ | $6.9 \times 10^{-4}$ | 4.6 | 3 | 0.6 |
| T110A | $4.5 \times 10^5$ | $1.9 \times 10^{-4}$ | 0.4 | 0.3 | -0.8 |
| S112A | $2.1 \times 10^5$ | $5.4 \times 10^{-4}$ | 2.5 | 2 | 0.3 |

**Supplementary Data Table 2.** Conserved scaffold residues of the NorM-Nb17_4, DMT-Nb16_4 and Lyso-Nb_4 in the NabFab-Nanobody interface interacting with a Heavy chain, b Light Chain of NabFab from PDB IDs: 7PHP, 7PIJ, 7RTH. The interface analysis was done in PISA(68). Chain “m” of the Nb and Chain “M/N”(Light chain/Heavy chain) of the Fab were considered for the interface analysis from the crystal structure 7RTH. The buried surface area is reported as the mean ± standard deviation calculated from the three structures

| Nb residues | Buried Surface Area (Å²) |
| --- | --- |
| Val 5 | 7.0 ± 3.6 |
| Glu 6 | 10.2 ± 3.4 |
| Gly 9 | 7.7 ± 2.4 |
| Gly 10 | 16.2 ± 3.7 |
| Leu 11 | 48.2 ± 8.9 |
| Gln 39 | 60.7 ± 7.6 |
| Ala 40 | 0.6 ± 0.1 |
| Pro 41 | 63.5 ± 6.7 |
| Gly 42 | 43.5 ± 17.3 |
| Lys 43 | 9.3 ± 2.2 |
| Glu 44 | 6.7 ± 1.9 |
| Arg 45 | 51.5 ± 17.4 |
| Thr 87 | 8.7 ± 0.9 |
| Ala 88 | 8.0 ± 1.9 |
| Val 89 | 39.8 ± 4.7 |
| Tyr 91 | 25.3 ± 7.6 |
| Trp 103 | 21.8 ± 6.5 |
| Gly 104 | 4.7 ± 1.6 |
| Lys 105 | 99.4 ± 3.7 |
| Gly 106 | 17.5 ± 2.9 |
| Thr107 | 2.3 ± 3.2 |
| Pro 108 | 79.7 ± 6.8 |
| Val 109 | 2.1 ± 0.9 |
| Thr 110 | 30.8 ± 4.0 |

**b**
| Nb residues | Buried Surface Area (Å²) |
| --- | --- |
| Pro 41 | 66.3 ± 3.7 |
| Gly 42 | 28.3 ± 4.8 |
| Lys 43 | 22.4 ± 4.6 |
| Thr 87 | 36.2 ± 3.6 |
| Thr 110 | 32.2 ± 2.1 |
| Val111 | 4.9 ± 2.0 |
| Ser 112 | 33.3 ± 1.1 |

**Supplementary Data Table 3.** Cryo-EM data collection, refinement, and validation statistics as defined by PHENIX(59).

| Electron cryomicroscopy data collection |  |  |  |  |
| --- | --- | --- | --- | --- |
| Sample | VcNorM NabFab complex |  | ScaDMT NabFab complex | ScaDMT NabFab complex dimer |
| PDB ID / EMDB ID | 7PHP / 13425 (Map1) / 13424 (Map2) |  | 7PIJ / 13438 | 7PHQ / 13426 |
| Microscope | FEI Titan Krios |  |  |  |
| Voltage (kV) | 300 |  |  |  |
| Camera | Gatan K3 |  |  |  |
| Energy filter | Gatan BioQuantum 1967 |  |  |  |
| Magnification (nominal) | 130,000 x |  |  |  |
| Pixel size (Å) | 0.67 |  | 0.66 |  |
| defocus range (µm) | -0.6 to -2.8 |  |  |  |
| 3D reconstruction |  |  |  |  |
| Map | Map1 | Map2 |  |  |
| Number of micrographs that went into processing | 9,711 | 9,711 | 17,957 |  |
| Final number of particles | 433,559 | 433,559 | 314,541 | 34,100 |
| Resolution (Å) | 3.68 | 3.47 | 3.78 | 8.45 |
| Resolution Range (Å) | 3.3-4.2 | 3.3-4.2 | 3.6-4.4 | n/a |
| FSC threshold | 0.143 | 0.143 | 0.143 | 0.143 |
| Map sharpening B factor | -151.127 | -109.28 | -152.2 | n/a |
| Coordinate refinement statistics |  |  |  |  |
| Initial model used (PDB ID) | 6ANI |  | 6ANI, 4WGV | 7PIJ (rigid body docking only) |
| Non-hydrogen atoms | 8647 |  | 8194 | 16388 |
| Protein residues | 1134 |  | 1068 | 2136 |
| RMSD bonds (Å) | 0.006 |  | 0.016 | 0.016 |
| RMSD angles (°) | 0.757 |  | 1.396 | 1.396 |
| Ramachandran favored (%) | 93.04 |  | 89.30 | 89.30 |
| Ramachandran allowed (%) | 6.96 |  | 10.61 | 10.61 |
| Ramachandran outliers (%) | 0.00 |  | 0.09 | 0.09 |
| Rotamer outliers (%) | 0.11 |  | 0.11 | 0.11 |
| Cβ outliers (%) | 0.00 |  | 0.00 | 0.00 |
| Clashscore | 6.49 |  | 13.08 | 13.08 |
| MolProbity score | 1.80 |  | 2.19 | 2.19 |

**Supplementary Data Table 4.** X-ray data collection, refinement, and validation statistics as defined by PHENIX(59).

|  |  |
| --- | --- |
| PDB ID | 7RTH |
| Crystal parameters: |  |
| Resolution (last shell) (Å) | 111.43-3.19 (3.36-3.19) |
| Space group | C121 |
| Unit cell parameters: |  |
| a, b, c (Å) | 225.53, 130.28, 223.98 |
| $\alpha$, $\beta$, $\gamma$ (°) | 90, 107.99, 90 |
| Matthews coefficient (Å<sup>3</sup>/Da) | 2.92 |
| Solvent content (%) | 57.53 |
| Data collection: |  |
| Completeness (%) | 99.4 (97.9) |
| No of unique reflections | 102005 |
| I/$\sigma$(I) | 9.5 (1.9) |
| R<sub>pim</sub> (%) | 17 (73.7) |
| CC (1/2) | 0.98 (0.47) |
| Redundancy | 5.7 (5.4) |
| Wilson B-factor (Å<sup>2</sup>) | 64.2 |
| Refinement: |  |
| R (%) / R<sub>free</sub> (%) | 21.6 / 26.7 |
| r.m.s.d. bond length (Å) | 0.002 |
| r.m.s.d. bond angle (°) | 0.445 |
| Average B value (Å<sup>2</sup>) | 77 |
| Number of molecules in AU | 9 |
| No of atoms: |  |
| Protein | 37949 |
| Water molecules | 165 |
| Other solvents | 354 |
| MolProbity analysis: |  |
| Ramachandran favored (%) | 95.4 |
| Ramachandran outliers (%) | 0.1 |
| Rotamer outliers (%) | 0.7 |
| C-beta outliers | 0 |
| Clashscore | 4.66 |
| Overall score | 1.56 |

r.m.s.d., root-mean-square deviation; AU, asymmetric unit.

